# A method for simultaneously monitoring phloem and xylem reconnections in grafted watermelon seedlings

**DOI:** 10.1101/2021.08.07.455541

**Authors:** Jianuo Xu, Xiaoyang Wei, Mu Xiong, Ting Zhang, Changjin Liu, Zhilong Bie, Yuan Huang

**Affiliations:** Key Laboratory of Horticultural Plant Biology, Ministry of Education/College of Horticulture and Forestry Sciences, Huazhong Agricultural University, Wuhan 430070, P.R. China; Centre for Plant Science, School of Environmental and Life Sciences, The University of Newcastle, Callahan NSW 2308, Australia; Agricultural Genomics Institute at Shenzhen, Chinese Academy of Agricultural Sciences, Shenzhen Branch, Guangdong Laboratory for Lingnan Modern Agriculture, Genome Analysis Laboratory of the Ministry of Agriculture, Guangdong, Shenzhen 518000, P.R. China

**Keywords:** Esculin, Acid fuchsin, Vascular reconnection, Watermelon, Cucurbit species, Grafting

## Abstract

Grafting is an effective way to increase watermelon tolerance to biotic and abiotic stresses. However, the survival of grafted seedlings largely depends on successful graft formation. Therefore, understanding the graft formation process, particularly the vascular reconnection process is of critical importance. This study found that lignin in watermelon stem shows strong auto-fluorescence under blue-light excitation which makes blue-light excited fluorescent tracers (FTs) such as 5(6)-carboxy fluorescein diacetate (CFDA) become unsuitable for assaying vascular connectivity in watermelon. In contrast, UV-light excited esculin and red-light excited acid fuchsin were proved to be efficient FTs for monitoring the phloem and xylem connectivity, respectively, in self-grafted watermelon. Furthermore, a combined application of esculin to the scion cotyledon and acid fuchsin to the rootstock root enabled simultaneous monitoring of the phloem and xylem connectivity in individual self-grafted watermelon seedlings. In addition, this method is also applicable in investigating the phloem and xylem reconnections in self-grafted melon and cucumber, and heterograft of watermelon, melon and cucumber onto pumpkin rootstock. Based on this established method, we found that phloem and xylem reconnections are not timely separated in self-grafted watermelon. Furthermore, low temperature and removal of the rootstock cotyledons both delayed the vascular reconnection process in watermelon. In conclusion, this new method provides a convenient, accurate and rapid way to analyze the vascular connectivity not only in watermelon, but also in other cucurbit crops.

## 1. Introduction

Vegetable grafting is extensively used today in agricultural production to control soil-borne pathogens (Yetisir et al., 2003; Thies et al., 2016), tolerance to salinity (Yang et al., 2013), suboptimal temperatures (Li et al., 2014; Yang et al., 2016), and mineral deficiency (Nawaz et al., 2017). Commercial vegetable grafting is mainly practiced in Cucurbitaceae and Solanaceae species, among them watermelon has the highest grafting proportion, more than 40% of watermelon plants are grafted in China (Zhong et al., 2018). Although grafting plays a central role in successful production of vegetables, however, the survival of grafted seedlings largely depends on successful graft formation. Therefore, understanding the graft formation process, particularly the phloem and xylem reconnection process is of critical importance.

Fluorescent tracers (FT) with a range of spectral properties have been used to assay phloem and xylem connectivity in various plants (Yin et al., 2012; Melnyk et al., 2015; Jiang et al., 2019; Tsutsui et al., 2020; Cui et al., 2021; Miao et al., 2021; Deng et al., 2021). Generally, a FT is loaded into a cut in the target tissue and the sequential dispersion of the reporter into other parts of the plant demonstrates the phloem and/or xylem reconnection. 5(6)-carboxy fluorescein diacetate (CFDA) perhaps is the most widely used FT in the plant. CFDA is non-fluorescent but it could be converted into fluorescent carboxy fluorescein (CF) by intracellular esterases in live cells and then be loaded into vascular for long distant transport. By applying CFDA to the self-grafted Arabidopsis, Melnyk et al. (2015) successfully documented the reconnecting process of vascular tissues and found that the phloem and xylem reconnections are temperately separated. However, resolution of current monitoring methods including the CFDA application applied by Melnyk et al. (2015) is limited. To our knowledge, current methods using FTs to assay the vascular connectivity are only able to conduct the phloem and xylem assays separately in the plant, making monitoring the phloem and xylem connectivity in individual plants impossible.

Furthermore, application of FT is often limited by endogenous autofluorecence in the plant, which is an ever-present roadblock for researchers to visualize specific fluorescent markers such as CFDA (Billinton and Knight, 2001; Donaldson, 2020). Common sources of this autofluorescence in the plant include flavins, NADH and NADPH, elastin and collagen, lipofuscins and lignin, and chlorophyll (Billinton and Knight, 2001). The presence of autofluorescence interferes with the FT which decreases contrast and clarity in fluorescence microscope visualization, and thus makes the FT application ineffective. In addition, autofluorecence spectra are generally broad. Hence, clarifying the autofluorescence source is important which provides the base for the selection of other applicable FTs.

Esculin and acid fuchsin are potential FTs to simultaneously monitor the phloem and xylem reconnections in grafted plants (Yin et al., 2012; Knoblauch et al., 2015; Miao et al., 2021; Deng et al., 2021). However, whether esculin is applicable in watermelon remains unknown. Esculin is a fluorescent coumarin glucoside which could be loaded into the phloem for long distance transport by the sucrose transporter AtSUC2 in Arabidopsis (Knoblauch et al., 2015; Knox et al., 2018). However, the barley sucrose transporter, HvSUT1, expressed in Arabidopsis failed to load esculin into the phloem (Knoblauch et al., 2015), putting a question mark on application of esculin in other species. Whereas, acid fuchsin has been used to monitor the xylem connectivity in Arabidopsis (Flaishman et al., 2008; Yin et al., 2012) and in cucumber (Miao et al., 2021). However, to our knowledge, acid fuchsin has only used as a mobile dye rather than being regarded as FT in these studies, making the application of this fluorescent molecule less sensitive.

In this study, we first elucidated that lignin in the watermelon stem shows strong autofluorescence under blue-light excitation, suggesting blue-light excited FTs such as CFDA is unsuitable for being used as FTs for assaying vascular connectivity in grafted watermelon. Base on this finding, we established that esculin and acid fuchsin are suitable for monitoring the phloem and xylem connectivity in grafted watermelon, respectively. Furthermore, a combined application method was developed in self-grafted watermelon seedlings to achieve the simultaneously monitoring of phloem and xylem reconnections in individual plants. Based on this method, we confirmed that effects of temperature and rootstock cotyledon on vascular reconnection in grafted watermelon are significant. The method introduced in this study provides an accurate, continent and inexpensive way to analyze the vascular connectivity in grafted watermelon.

## 2. Materials and methods

### 2.1. Plant material and cultivation

The experiment was conducted in plant growth room at the National Center of Vegetable Improvement in Huazhong Agricultural University, Central China (latitude, 30° 27’ N; longitude, 114° 20’ E). Watermelon cultivar 97103 (*Citrullus lanatus*, Beijing Vegetable Research Center, Beijing Academy of Agriculture and Forestry Sciences, China), melon cultivar Akekekouqi (*Cucumis melo*, Hami Melon Research Center, Xinjiang Academy of Agricultural Sciences, China), cucumber cultivar Jinyou35 (*Cucumis sativus*, Cucumber Research Institute of Tianjin Kernel Agricultural Science and Technology Company Limited, China), and interspecific pumpkin hybrid cultivar Qingyanzhen No.1 (*C. maxima* × *C. moschata*, Qingdao Academy of Agricultural Sciences) were used as the plant materials in this study. Seeds were sterilized for 15 min with 0.1% KMnO_4_ and soaked in water at 55 °C for 20 min. Then, seeds were transferred onto wet filter papers and cultured in petri dishes at 30 °C in darkness for 36 h. Germinated seeds were then sown in 72- cell plug tray (540 mm × 280 mm) in mixed media (peat: perlite: vermiculite, 7:3:1, volume ratio), and grown in the plant growth room. During the cultivation, a standard growth condition was maintained with 150 μmol·m^-2^·s^-1^ of photosynthetic photon flux density (PPFD), 14/10 h photoperiod, day/night temperature at 28°C/18°C, relative humidity at 65-85%.

### 2.2. Grafting and healing

In grafting, 11- and 8-day-old watermelon seedlings were used as the rootstock and the scion, respectively. One cotyledon grafting was performed as described by Hassell et al. (2008). Six graft combinations were used in this study including self-grafted watermelon, melon and cucumber, and three heterograft combinations that were watermelon, melon and cucumber grafted onto pumpkin, respectively. For rootstock cotyledon removal experiment, splice grafting was conducted as described by Devi et al. (2021). The plants were placed in the healing chamber after grafting and this day was defined as 0 day after grafting (DAG).

The healing conditions were: from 0 to 1 DAG, maintained in dark; from 1 to 6 DAG, grown under low light condition (50 μmol·m^-2^·s^-1^, 14/10 h photoperiod); from 7 to 9 DAG, normal light condition (150 μmol·m^-2^·s^-1^, 14/10 h photoperiod). The day and night temperatures in healing chamber were 26°C. The humidity was kept above 95% during the first 5 days (0 to 4 DAG), then decreased to about 75% and maintained at a period from 5 to 9 DAG. At 10 DAG, plants were transferred from the healing chambers to the growth room and grown under standard condition as described in section 2.1.

### 2.3. Paraffin sectioning, staining and confocal imaging of watermelon graft junction

Graft junctions were sampled from self-grafted watermelon seedlings at 6 DAG. To label lignified tissues in stem sections, samples were stained with 1% (w/v) safranin for 2 h. After staining, samples were mounted in 50% glycerol and imaged using a confocal laser scanning microscope (Leica SP8, Leica, Germany). For safranin, fluorescence was detected at 552 nm excitation, 560-650 nm emission wavelength; for auto-fluorescence, fluorescence was detected at 488 nm excitation, 490-530 nm emission wavelength.

### 2.4. Phloem and xylem connectivity assays simultaneously

Phloem and xylem connectivity were measured simultaneously using the same plant by esculin (E8250, Sigma-Aldrich) and acid fuchsin (F8129, Sigma-Aldrich) movement across the graft junction, respectively. The value was presented as (reconnected plants/total plants) × 100%. Before the esculin and acid fuchsin applications, the adaxial surface of the scion cotyledon was rubbed gently with sandpaper to remove the wax, whereas the root was cut off to 3-5 cm long. Afterwards, 2% (w/v) esculin dissolved in 60% (v/v) acetonitrile together with 2.5 mM ethylene diamine tetra-acetic acid (EDTA) were applied to the scion cotyledon to measure phloem connectivity. Simultaneously, 0.5% (w/v) acid fuchsin was applied to the root to measure xylem connectivity, using the same plant for phloem connectivity measurement. Grafted seedlings were then kept in dark at room temperature for 2 h. Stem sections from the scion or rootstock 0.5 cm above or below the graft junction, respectively, were sampled for fluorescence assay. For esculin, a standard filter set for UV was applied which includes a 360/40 nm excitation filter and a 420 LP barrier filter. For acid fuchsin, a standard filter set for DSRed was applied which includes a 545/30 nm excitation filter and a 620/60 nm barrier filter.

### 2.5. Splice grafting treatment

The self-grafted watermelon seedlings were used as materials. There were two treatments in this experiment, i.e., (1) one cotyledon grafting with one rootstock cotyledon, (2) splice grafting without rootstock cotyledon. Phloem and xylem reconnections were measured at day 3, 6 and 9 DAG.

### 2.6. Suboptimal low temperature treatment

The self-grafted watermelon seedlings were used as materials, grafted plants were healed at 26°C or 18°C (suboptimal low temperature), the other conditions were the same as described in above 2.2. Phloem and xylem reconnection were measured at day 3, 6 and 9 DAG.

### 2.7. Statistical analysis

Statistical analysis was performed using SPSS 25.0 software (SPSS Inc., Chicago, IL, USA). The data were presented as means ± SE of three replicates. Each replicate had 20 plants. Student’s t-test was conducted to show the difference of phloem and xylem reconnections between 26°C and 18°C healing conditions, and between one cotyledon grafted and splice grafted plants. Levels of significance were represented by **p < 0.01 or ***p < 0.001.

## 3. Results

### 3.1. Lignin in watermelon stem tissues showed strong green auto-fluorescence under the blue-light excitation

By observing stem cross sections of watermelon stems using a fluorescence stereomicroscope, we found that watermelon stem showed strong green auto-fluorescence under blue-light excitation (a standard filter set for GFP viewing) (Figure 1A-C). Hence, to clarify the source of this green autofluorescence in watermelon, we then performed safranine staining on stem sections. Safranine is a fluorescent dye commonly used for labelling lignified tissue in plants (Bond et al., 2008). As shown in Figure 1 D-G, the green auto-fluorescence in watermelon stem co-localized with the safranine stained lignified tissues including the xylem and the stem bark, indicating the green auto-fluorescence source was lignin. In addition, from our observation, other agricultural cucurbits crops including melon and pumpkin stems also showed strong autofluorescence under blue-light excitation (Supplemental Figure S1).

**Figure 1.**
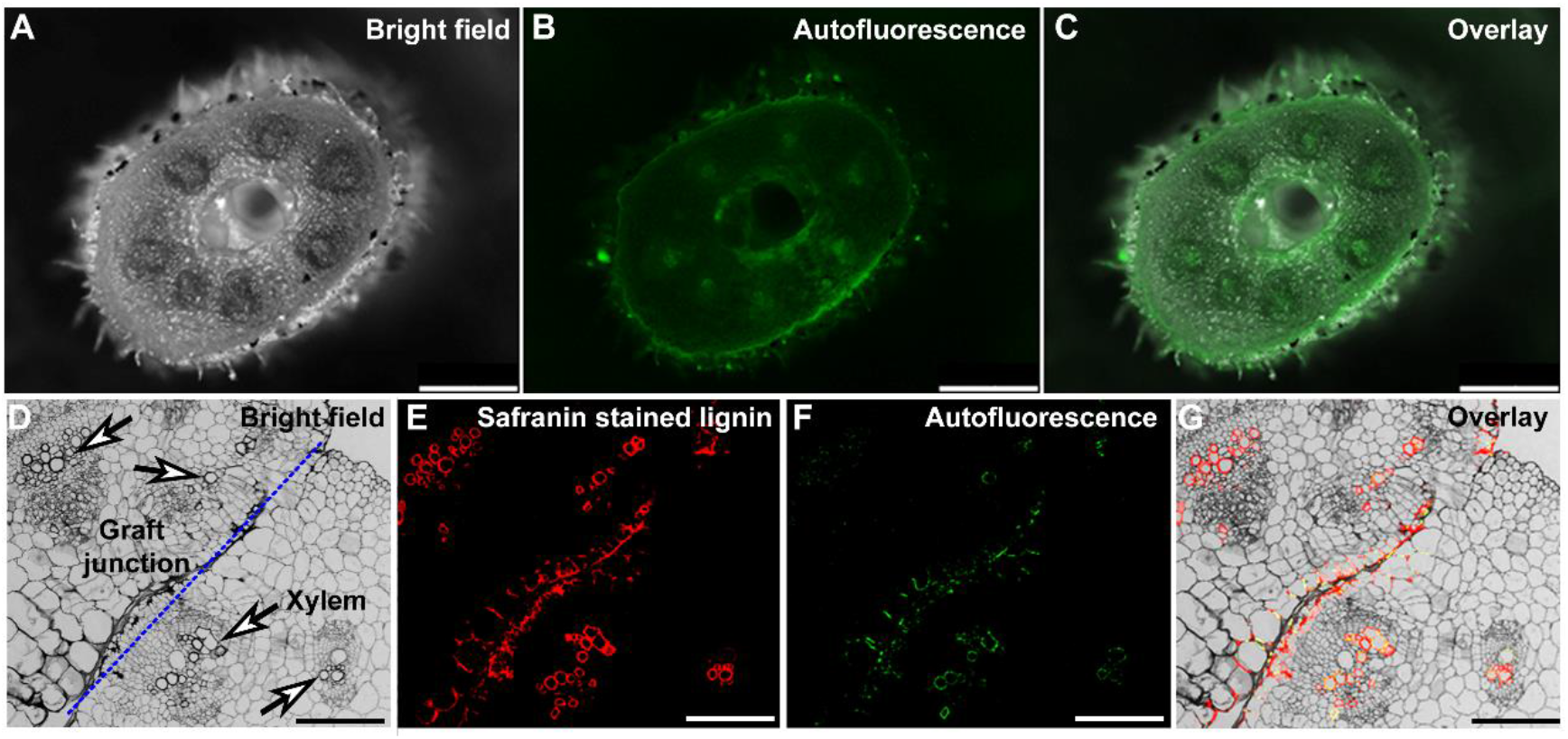
Watermelon stem shows strong lignin induced auto-fluorescence under blue light excitation. The hand-cutting stem sections of 10-day-old watermelon seedlings were imaged and showed strong auto-fluorescence when being excited by the laser for GFP detection. (A-G) Images of a cross section of watermelon stem viewed by a fluorescence stereomicroscope, (A) Bright field; (B) Auto-fluorescence under blue-light excitation; (C) Overlay image of (A) and (B); (D-G) Images of a paraffin cross section of watermelon graft junction viewed by a confocal microscope, (D) Bright field; (E) Lignin staining with safranin; (F) Auto-fluorescence of the graft junction under blue-light excitation; (G) Overlay image of (D-F). Dash line indicates graft junction. Arrows indicate xylem tissues. Bar in (A-C): 1 mm; bar in (D-G): 0.2 mm.

### 3.2. A modified technique using mobile FTs to simultaneously monitor phloem and xylem connectivity in grafted watermelon seedlings

To avoid noise from the lignin auto-fluorescence in watermelon stem, we selected two fluorescent molecules esculin and acid fuchsin that are not excited by blue-light and tested their application in watermelon. The absorbance/emission wavelengths for esculin and acid fuchsin are 367/454 nm and 540/630 nm (Supplemental Table S1), respectively. Thus, theoretically, fluorescent signals of these two fluorescent molecules would not be interfered by tissue auto-fluorescence in watermelon stem, nor have interference with each other. Indeed, under the stereomicroscope used in this study, esculin and acid fuchsin were excited only by the UV-light and the red-light, respectively. In addition, watermelon stem did not show discernible auto-fluorescence under UV-light (for esculin excitation) nor red-light (for acid fuchsin excitation) excitations. Hence, we reasoned that esculin and acid fuchsin might be suitable to be used as FTs in watermelon. Furthermore, applying esculin and acid fuchsin to an individual grafted seedling at the same time might be able to simultaneously monitor the phloem and xylem connectivity in the plant.

Therefore, to test the applicability of esculin in watermelon, and to further test the possibility of achieving a simultaneously monitoring of the phloem and xylem connectivity in grafted watermelon, a combined application with esculin and acid fuchsin was applied in survived watermelon seedlings at 12 DAG. As shown in Figure 2A, 2% (w/v) esculin and 0.5% (w/v) acid fuchsin were applied to the scion cotyledon and the rootstock root of survived grafted watermelon seedlings, respectively. Cross sections of the scion and the rootstock stems 0.5 cm away from the graft junction were sampled 2 h after the treatment and the fluorescent signals were detected using a fluorescent stereomicroscope. As shown in Figure 2, esculin applied onto the scion cotyledon had been loaded into the scion phloem (Figure 2B) and then been transported to the rootstock (Figure 2D) through the graft junction via reconnected phloem. Meanwhile, acid fuchsin fed to the rootstock root had been loaded into the rootstock xylem (Figure 2E) and transported to the scion (Figure 2C) through the reconnected xylem. Together, esculin and acid fuchsin could be applied as FTs to monitor the phloem and xylem connectivity in grafted watermelon.

**Figure 2.**
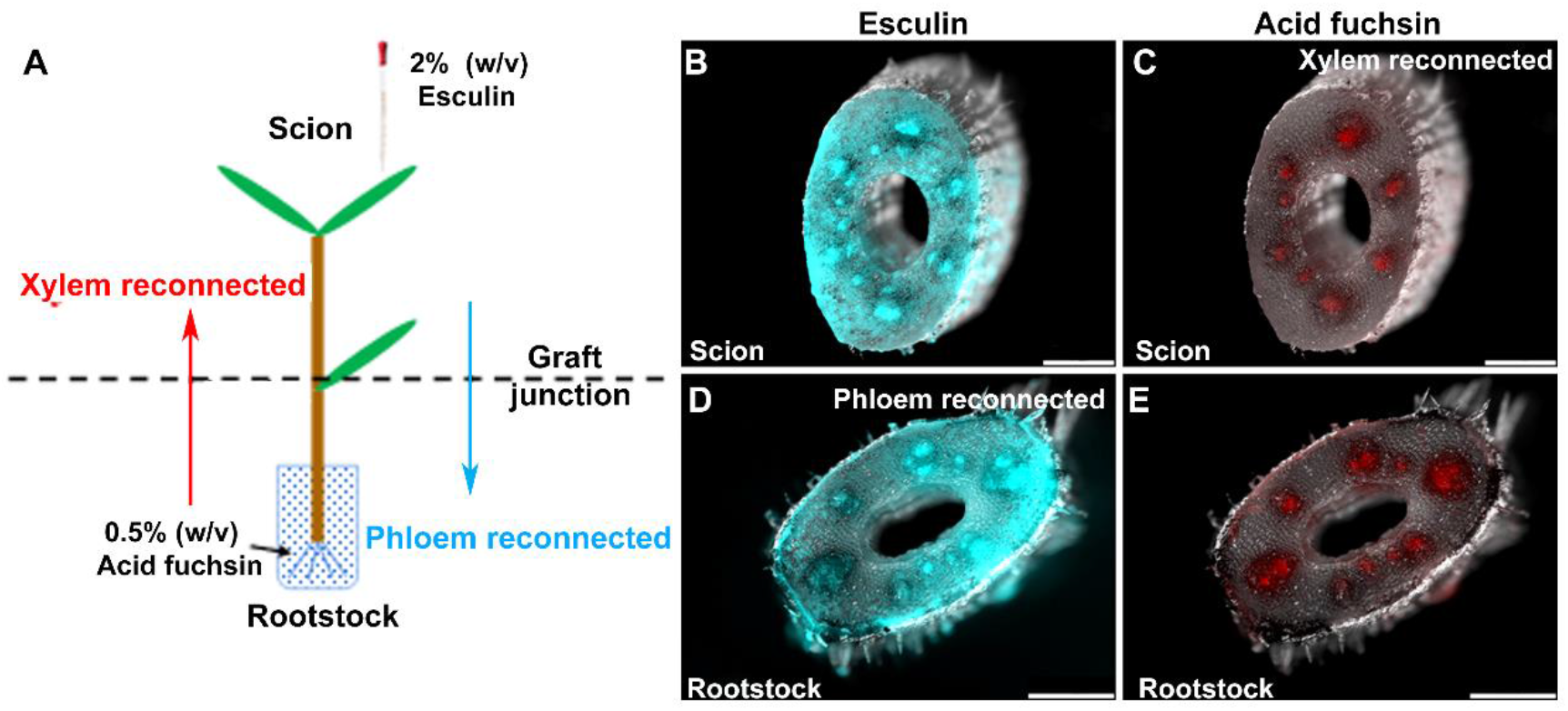
Application of esculin and acid fuchsin in simultaneously monitoring the phloem and xylem connectivity in grafted watermelon. Survived grafted watermelon seedlings at 12 DAG were used in this test. Fluorescent molecules esculin and acid fuchsin were applied to scion cotyledon and rootstock root, respectively, to test their application in monitoring the phloem and xylem connectivity in grafted watermelon. (**A**) A schematic diagram illustrates the esculin and acid fuchsin treatments to grafted watermelon, as well as their transport route in the plant. (**B**) Esculin detected in the scion stem. (**C**) Acid fuchsin detected in the scion stem. (**D**) Esculin detected in the rootstock stem; (**E**) Acid fuchsin detected in the rootstock stem. Arrows in (**A**) indicate transport directions: blue, esculin from scion to rootstock; red, acid fuchsin from rootstock to scion. Bar: 1 mm.

Furthermore, the mobility of esculin and acid fuchsin in phloem and xylem, respectively, are comparable to one another. As described above, after 2 h, exogenously applied esculin and acid fuchsin both had been transported through the grafted junction to their targeted sink tissues, respectively (Figure 2). Most importantly, as shown in Figure 2, fluorescence of these two molecules in the same watermelon stem showed no interference with each other. This esculin/acid fuchsin combined application is able to achieve a simultaneously monitor of the phloem and xylem reconnections in grafted watermelon seedlings.

### 3.3. Vascular reconnection in self-grafted watermelon seedlings

To investigate the vascular reconnecting process in self-grafted watermelon, we applied the combined esculin/acid-fuchsin. Phloem and xylem reconnections reached to 50% both at 5-6 DAG and to 100% both at 12-13 DAG (Figure 3A and B). In addition, during the reconnection process, phloem reconnection and xylem reconnection in individual seedlings occurred in a random order (Figure 3C). For instance, we observed four different types regarding phloem and xylem reconnections at 6 DAG, as indicated by the presences of esculin an acid fuchsin fluorescence in scion and rootstock stems, (i) Phloem and xylem both remain unconnected (Figure 4A-D, Supplemental Figure S2A-D); (ii) Phloem reconnected but xylem-unconnected (Figure 4E-H, Supplemental Figure S2E-H); (iii) Xylem reconnected but phloem un-connected (Figure 4I-L, Supplemental Figure S2I-L); (iv) Phloem and xylem both reconnected (Figure 4M-P, Supplemental Figure S2M-P). Statistical analysis demonstrated that, in 6 DAG grafted seedlings, around 32% seedlings only had reconnected phloem, 38% only had reconnected xylem, and 5% seedlings had both of the reconnected phloem and xylem (Figure 3C).

**Figure 3.**
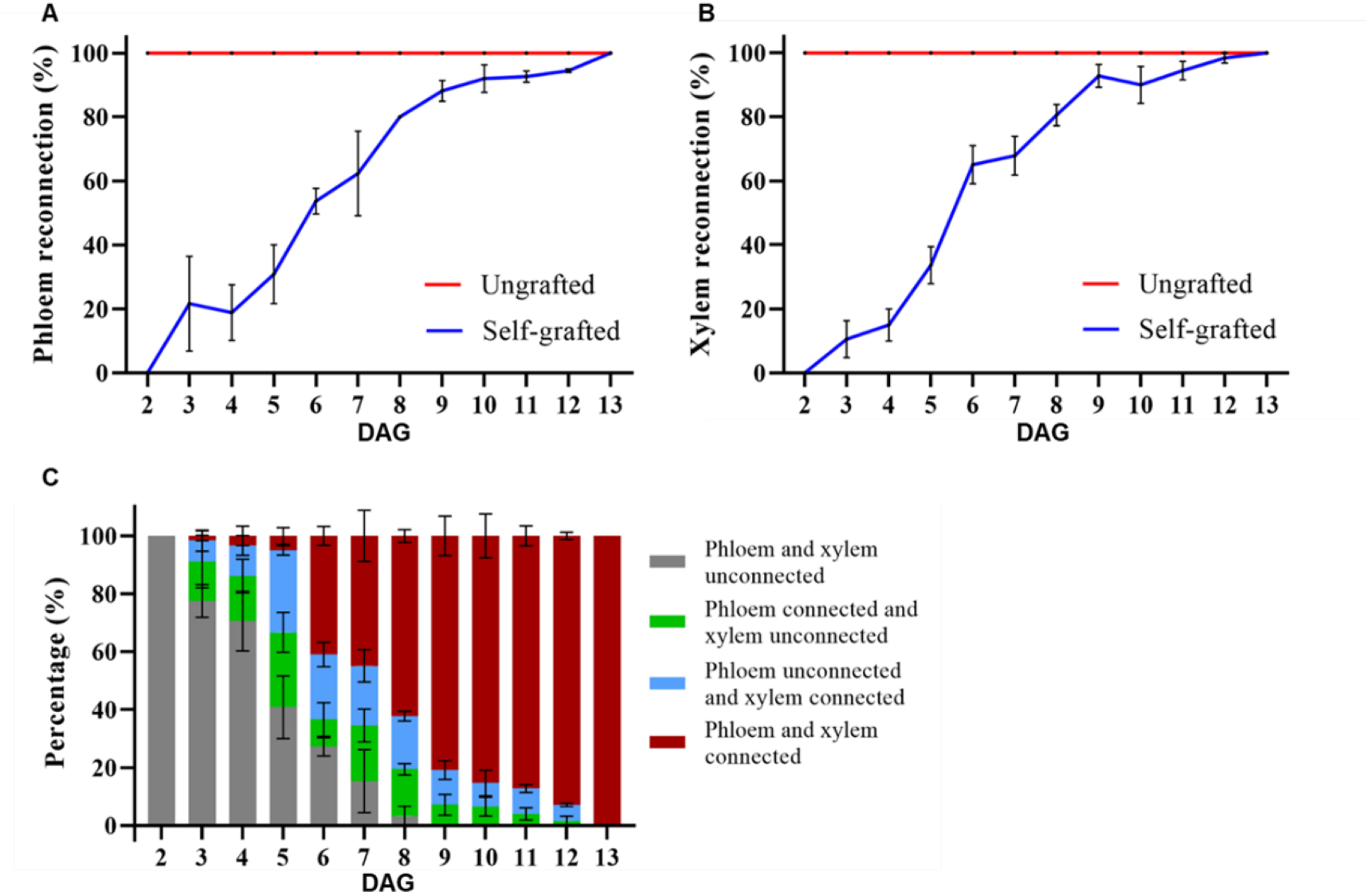
Phloem and xylem reconnections are not timely separated in self-grafted watermelon. Un-grafted seedlings were used as positive control. (**A**) Phloem and (**B**) xylem reconnections in grafted watermelon seedlings were similar to each other. (**C**) Percentage of four different vascular reconnection types observed in self-grafted seedlings during graft formation. Data of each time point was collected from three independent experiments with 20 seedlings per experiment. DAG: days after grafting.

**Figure 4.**
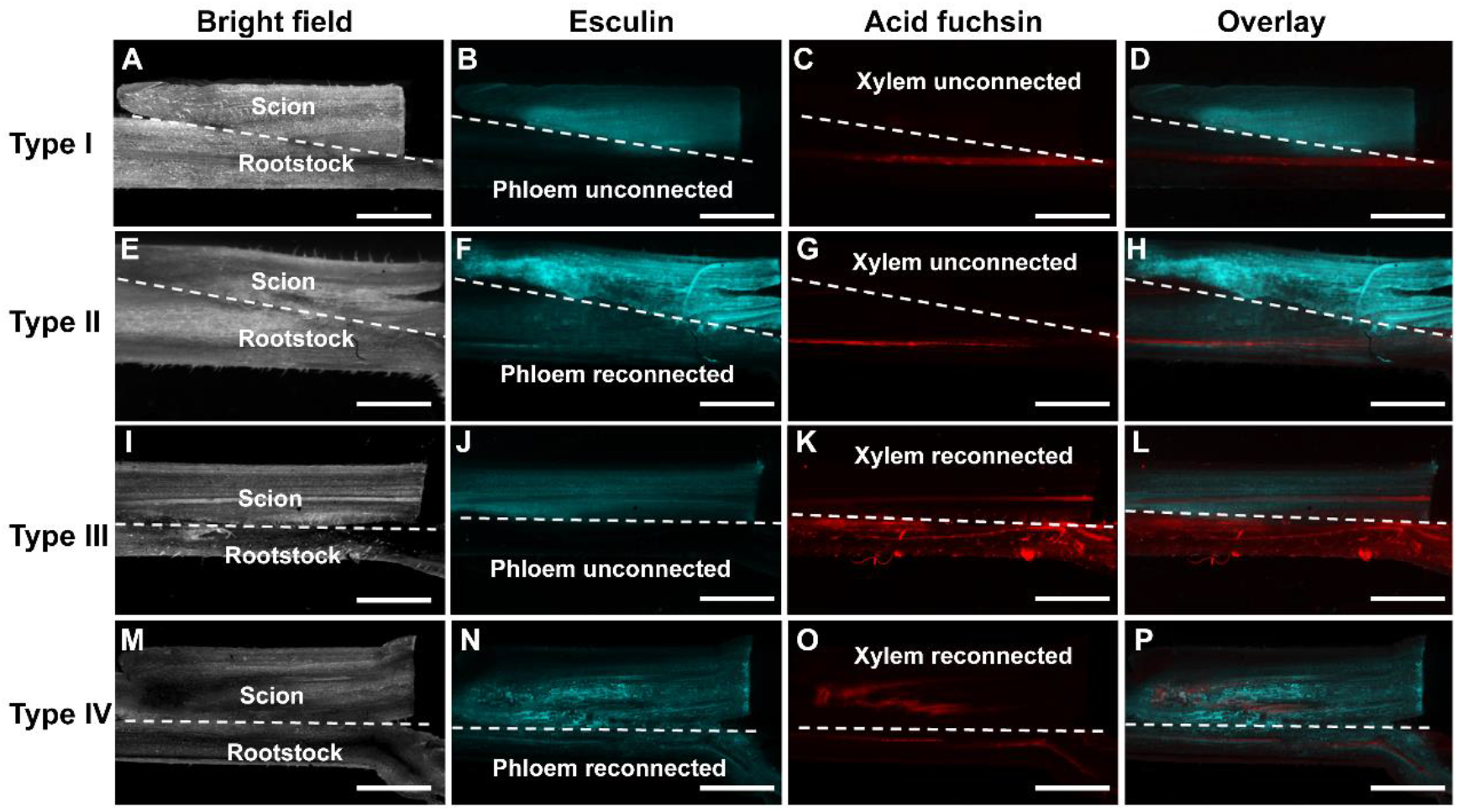
Four different reconnection types in self-grafted watermelon seedlings at day 6 after grafting (longitudinal cutting). To monitor phloem and xylem reconnections in grafted watermelon seedlings, fluorescent tracers including esculin and acid fuchsin were applied on the scion cotyledons and the rootstock roots, respectively. Stem sections from the scion and the rootstock of the same grafted seedlings were examined for the presences of esculin (Cyan) and acid fuchsin (Red) using fluorescent dissecting microscope. (A-D) Type I, grafted seedling with un-reconnected phloem and xylem. (E-H) Type II, grafted seedling with reconnected phloem and unconnected xylem. (I-L) Type III, grafted seedling with reconnected xylem and unconnected phloem. (M-P) Type IV, grafted seedling with reconnected phloem and xylem. Dash lines indicate graft junction. Bar: 2 mm.

### 3.4. The developed method demonstrates the impact of temperature on the vascular reconnection in self-grafted watermelon seedlings

Grafted watermelon seedlings were grown under 18 °C and 26 °C, respectively. Phloem and xylem reconnections were measures at 3, 6, and 9 DAG grafted seedlings. We observed that both of the phloem and xylem reconnections were significantly delayed in seedlings grown under 18 °C compared to 26 °C (Figure 5). Under 26 °C, reconnection rates of the phloem were 18% at 3 DAG, 58% at 6 DAG, and 80% at 9 DAG (Figure 5A). In contrast, under 18 °C, the phloem reconnection rates were 0% at 3 DAG, 20% at 6 DAG, and 50% at 9 DAG (Figure 5A). For xylem reconnection, at 3, 6, and 9 DAG, under 25°C, the rates were 18%, 61%, and 90%, respectively, whereas, under 18°C, the rates were decreased to 0%, 30%, and 61%, respectively. Together, this result confirms that low temperature delays vascular reconnection in watermelon grafted seedlings and in turn demonstrates that the combined application with esculin and acid fuchsin developed in this study is accurate and reliable.

**Figure 5.**
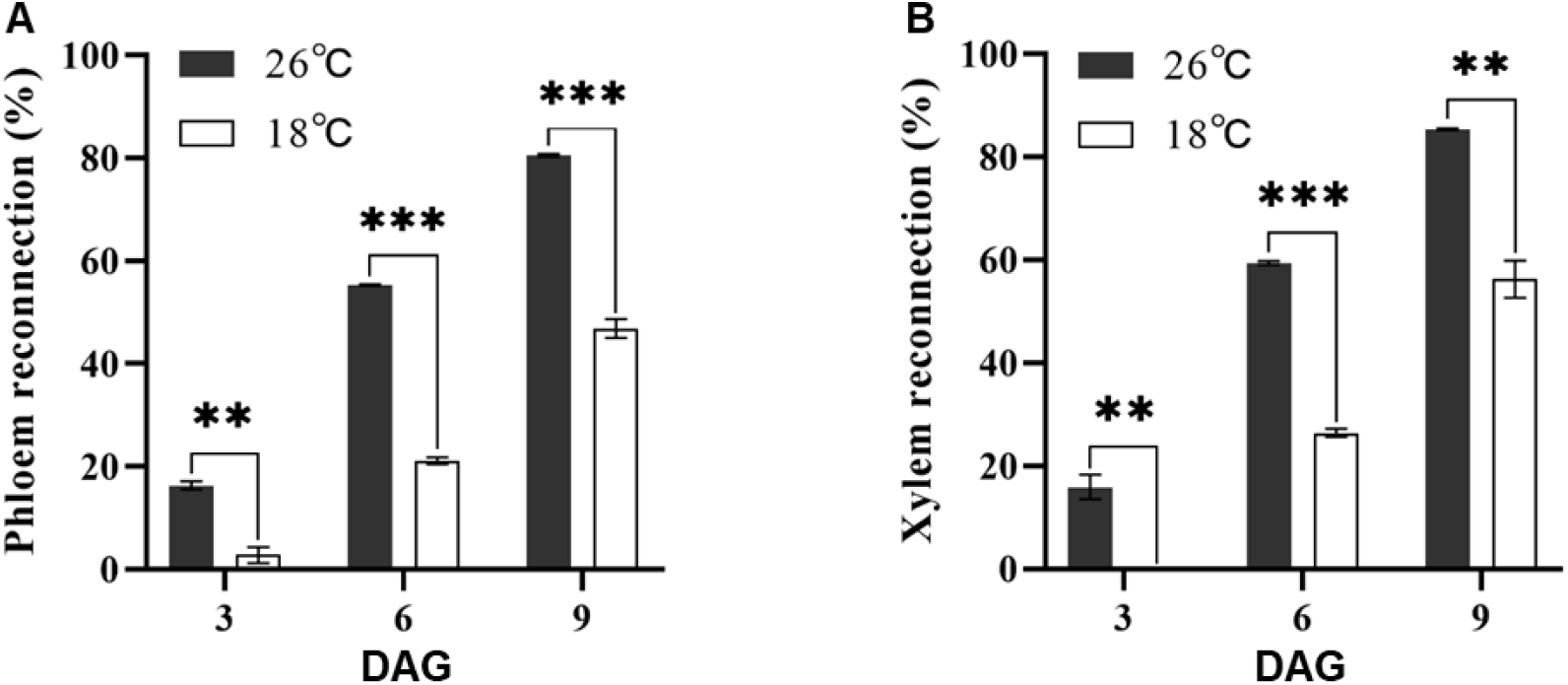
Suboptimal low temperature treatment delays both of the phloem and xylem reconnections in self-grafted watermelon seedlings. (A) Phloem reconnection rates. (B) Xylem reconnection rates. Asterisks indicate significant difference between 18°C and 26°C at 3 DAG, 6 DAG and 9 DAG, by Student’s t-test (\*\**p* < 0.01, \*\*\**p* < 0.001). DAG: days after grafting.

### 3.5. Rootstock cotyledon is essential for the vascular reconnection in self-grafted watermelon seedlings

To investigate the impact of rootstock cotyledon on vascular reconnection in grafted watermelon, we performed splice-grafting in watermelon seedlings and analyzed the vascular reconnection process using the esculin/acid fuchsin combined application. Based on results shown in Figure 6, reconnection rates of phloem and xylem both were significantly lower in splice-grated seedlings at 3, 6, and 9 DAG in comparison with control seedlings that have one rootstock cotyledon.

**Figure 6.**
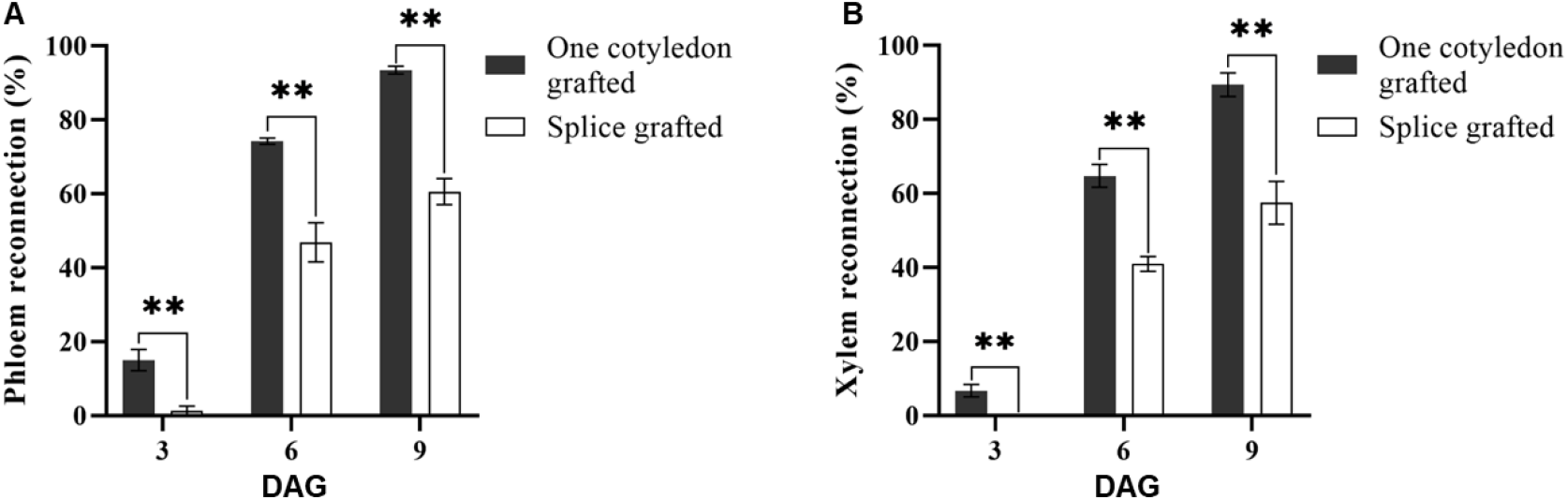
Rootstock cotyledon is essential for the phloem and xylem reconnections in self-grafted watermelon seedlings. (A) Phloem reconnection rate. (B) Xylem reconnection rate. One cotyledon grafted: grafted seedling with one rootstock cotyledon; Splice grafted: grafted seedling with no rootstock cotyledon. Asterisks indicate significant difference by Student’s t-test (** *p* < 0.01). DAG: days after grafting.

### 3.6. Application of the esculin/acid fuchsin combined treatment in other cucurbit species

To test general applicability of this method for monitoring the phloem and xylem connectivity in cucurbit species, we first applied the esculin/acid fuchsin combined application to self-grafted melon and cucumber seedlings. As shown in Figure 7, esculin applied to the scion cotyledons of self-grafted melon and cucumber was successfully loaded into the scion phloem (Figure 7B, H) and then transported into the rootstock (Figure 7E, K) through graft junctions of each species. Meanwhile, acid fuchsin fed to the rootstocks was detected in both of the scion and rootstock stems in self-grafted melon and cucumber seedlings (Figure 7C, F, I and L). Together, these results suggest that the esculin/acid fuchsin combined treatment is applicable in monitoring vascular connectivity in self-grafted melon and cucumber seedlings. Furthermore, we had also tested the applicability of this method in compatible heterograft combinations of watermelon, melon and cucumber scions onto pumpkin rootstock. As expected, esculin and acid fuchsin both could be loaded into and transported in vascular of heterograft seedlings of all three combinations (Supplemental Figure S3). These observations indicated the general applicability of this method for monitoring vascular connectivity in self-grafted and heterograft seedlings in cucurbit species.

**Figure 7.**
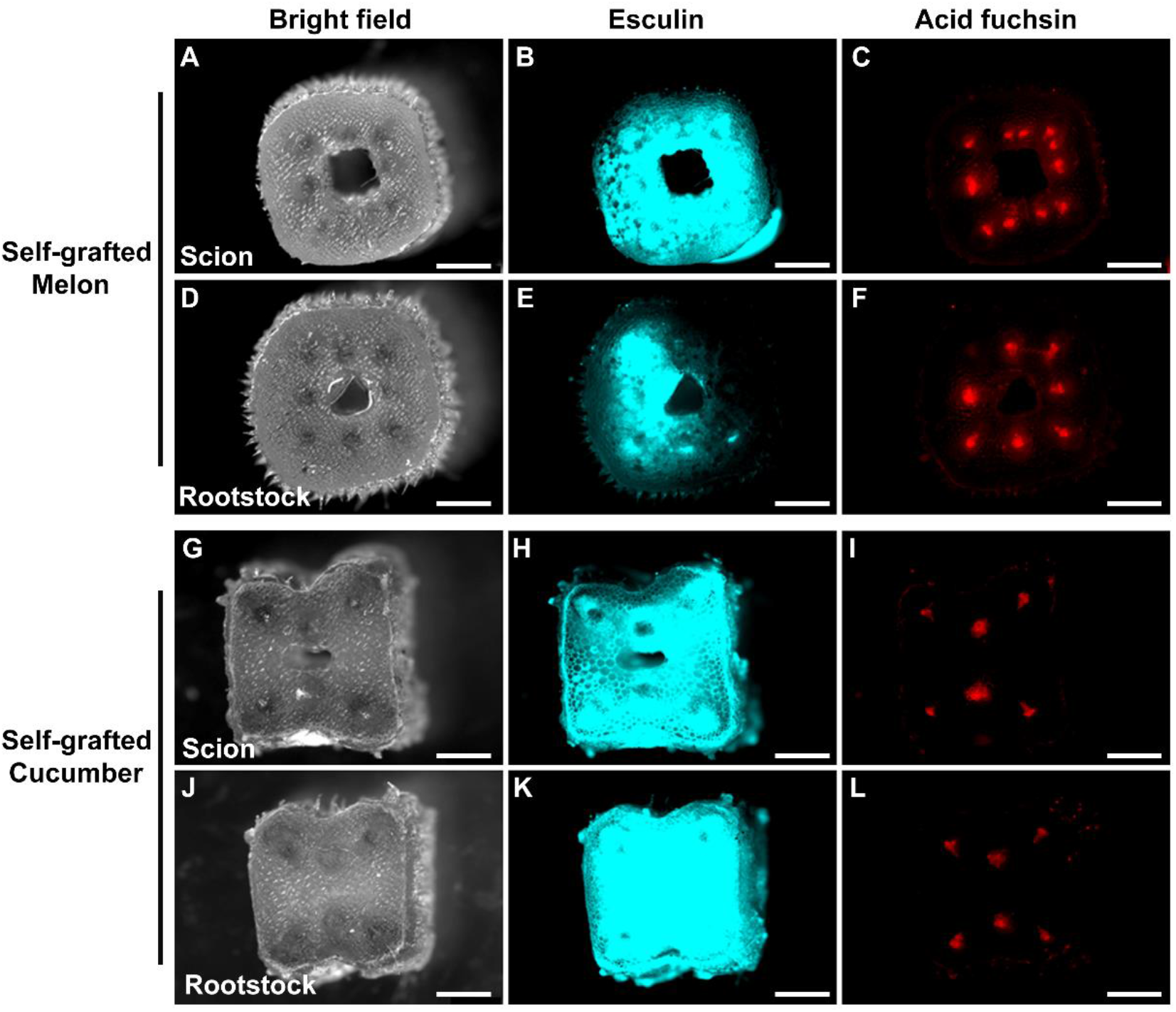
Application of esculin and acid fuchsin in self-grafted melon and cucumber seedlings. Self-grafted melon and cucumber seedlings at day 12 after grafting (DAG) were used in this test. (**A-F**) Self-grafted melon stem sections. (**G-L**) Self-grafted cucumber stem sections. Esculin fed on the scion cotyledon was successfully loaded into the scion phloem in both of the self-grafted (**B**) melon and (**H**) cucumber seedlings, and then was transported into the rootstock in both of self-grafted (**E**) melon and (**K**) cucumber seedlings through the graft junctions. Acid fuchsin fed on the rootstock root was loaded into the rootstock xylem in both of the self-grafted (**F**) melon and (**L**) cucumber seedlings, and then was transported into the scion in both of self-grafted (**C**) melon and cucumber seedlings. Bar: 1 mm.

## 4. Discussion

### 4.1. A combined application of esculin and acid fuchsin enabled simultaneous monitoring of the phloem and xylem connectivity

Fluorescent tracers (FTs), particularly carboxyfluorescein diacetate (CFDA), has been widely used to monitor symplastic transport through phloem and xylem in various plants (Oparka et al., 1994; Wright et al., 1996; Botha et al., 2008; Melnyk et al., 2015). However, its application is interfered by tissues autofluorescence in plants. In plants, one major green auto-fluorescent component (excitation at 488 nm) is lignin (Billinton and Knight, 2001; Donaldson, 2020). The peak absorbance/emission wavelengths of green lignin auto-fluorescence are 488/530 nm (Billinton and Knight, 2001; Donaldson, 2020). Consequently, the green lignin auto-fluorescence would interfere with blue-light-excited FTs, making the fluorescent signal unreliable. Therefore, blue-light-excited FTs, such as CFDA (peak excitation: 494 nm; peak emission: 521 nm) (Supplemental Figure S4), are not suitable for being used to assay vascular connectivity in watermelon (Figure 1).

Cucurbits crops including watermelon, melon and pumpkin show strong autofluorescence under the blue-light excitation, making blue-light excited FTs inefficient in monitoring the vascular connectivity in these species (Figure 1; Supplemental Figure S1, S4). Hence, we have developed a convenient, accurate and inexpensive esculin/acid fuchsin combined application method for simultaneously monitoring vascular connectivity in grafted seedlings in major agricultural cucurbits crops. The method consists of two independent applications, applying esculin to scion cotyledon to monitor phloem connectivity, meanwhile, applying acid fuchsin to rootstock root to determine xylem connectivity. Conventional monitoring methods used to apply only one type of FT such as CFDA to monitor the phloem or xylem connectivity in the plant (Melnyk et al., 2015; Jiang et al., 2019; Tsutsui et al., 2020; Cui et al., 2020; Miao et al., 2021; Yin et al., 2012; Deng et al., 2021). Consequently, the resolution of conventional methods in illustrating the vascular connectivity in individual plants is insufficient. This combined application of esculin and acid fuchsin in individual plants, however, is able to monitor the phloem and xylem connectivity in a single plant without the need to conduct the assays separately in two individuals, which greatly lower the workload and increases the depth of the determining method. The ability to analyze the phloem and xylem connectivity in individual plants enables the investigation of the vascular reconnection process at individual level in grafted plants, which makes the data more accurate. Furthermore, this method also provides a valid way to further investigate the vascular biology in the plant, particular to elucidate the development process of the phloem and xylem and their potential interactions during the process of graft formation in grafted plants. In addition, the success of this method in self-grafted watermelon, melon and cucumber seedlings and heterograft combinations of watermelon, melon and cucumber onto pumpkin demonstrates its general applicability in cucurbit crops.

### 4.2. Phloem and xylem reconnections are not timely separated in self-grafted watermelon

We used this method to analyze the vascular reconnection process in self-grafted watermelon. A previous study by Melnyk et al. (2015) described in grafted Arabidopsis seedlings, phloem and xylem reconnections are temporally separated that phloem reconnection is prior to xylem reconnection. Generally, phloem reconnection reached to 50% at 3 DAG and to 100% at 4 DAG, whereas these time points for xylem reconnection were at 6 DAG and 7 DAG, respectively (Melnyk et al., 2015). Recent study in the heterograft of cucumber onto pumpkin (Miao et al., 2021), however, showed slightly different results that the phloem and xylem reconnections in the heterograft seedlings are not totally timely separated but the phloem reconnection peak occurs two days earlier compared to the xylem. However, our observation in self-grafted watermelon indicated that the phloem and xylem reconnections are not timely separated that they occur at the same time (Figure 3). Nevertheless, we also observed that the phloem and xylem reconnections occur randomly in individual plants (Figures 3, 4), implying the mechanism regulating the vascular reconnection process is complex. This substantial difference might be due to difference in the plant species and grafting methods applied in different studies. For instance, we used one cotyledon grafting in watermelon, while Melnyk et al. (2015) used splice grafting without rootstock cotyledon in Arabidopsis. Besides, the limits of the monitoring method used in previous studies, which are only able to conduct the phloem and xylem connectivity assay in different seedlings, might cause evitable errors.

### 4.3. Application of the developed method to demonstrate the effects of temperature and rootstock cotyledon on the vascular reconnection of grafted watermelon

Previous study by Yang et al. (2016) indicated that low temperature inhibits vascular reconnection in plants, and the survival rate of the splice grafting watermelon seedlings that have no rootstock cotyledon is significantly lower compared with seedlings that have at least one rootstock cotyledon (Dabirian et al., 2017; Devi et al., 2021). However, the effects of temperature and the rootstock cotyledon on the vascular reconnection remains unclear. Using the developed method, we found that low temperature and rootstock cotyledon removal significantly delays the reconnection of phloem and xylem, which strongly supports the phenomenon that low temperature and rootstock cotyledon removal results in lower graft survival and seedling growth (Yang et al., 2016; Dabirian et al., 2017; Devi et al., 2021).

## 5. Conclusion

We have demonstrated that blue-light excited FTs are not suitable for watermelon, melon and pumpkin in their vascular connectivity assays. The method developed in this study for monitoring the vascular connectivity in grafted plants using a combined application with esculin and acid fuchsin has overcome this obstacle in cucurbit crops. This rapid method was used to monitor the phloem and xylem reconnection processes in self-grafted seedlings and found that the vascular reconnection process in watermelon is different from previous observation in other species. Temperature and the rootstock cotyledon both have significant effects on the vascular reconnection in watermelon. The ability of monitoring the phloem and xylem connectivity in individual plants greatly lowers the workload and enables future studies to investigate the vascular development process in plants more accurate and efficient.

## Acknowledgements

This research was funded by the National Natural Science Foundation of China (31972434), National Key Research and Development Program of China (2019YFD1001900), China Agriculture Research System of MOF and MORA (CARS-25), Hubei Provincial Natural Science Foundation of China (2019CFA017), and Huazhong Agricultural University-Agricultural Genomics Institute at Shenzhen, Chinese Academy of Agricultural Sciences Cooperation Fund (SZYJY2021005).

## Author contributions

YH, XYW and ZLB devised the project; JNX, XYW, TZ, MX and CJL performed experimental analyses; JNX, XYW, and YH performed data analyses. YH, XYW and JNX wrote the manuscript and all authors approved the final text.

