## Supplemental table and figures for "A method for simultaneously monitoring phloem and xylem reconnections in grafted watermelon seedlings"

### Supplementary materials

**Supplemental Table 1. Fluorescent tracers used in this study.**

| Fluorescent tracers | Molecular weight (g/mol) | Formula | λEx (nm) | λEm (nm) | Phloem transport | Xylem transport | Reference |
|---|---|---|---|---|---|---|---|
| **Esculin (6,7-dihydroxy-coumarin 6-b-D-glucopyranoside)** | 340.3 | $C_{15}H_{16}O_9$ | 367 | 454 | √ | √ | Knoblauch et al., 2015 |
| **Acid fuchsin** | 585.5 | $C_{20}H_{17}N_3Na_2O_9S_3$ | 540 | 630 | √ | √ | Yin et al., 2012 |
| **CFDA [5(6)-Carboxyfluorescein diacetate]** | 460.4 | $C_{25}H_{16}O_9$ | 454 | 521 | √ | √ | Melnyk et al., 2015 |


1. Knoblauch, M., Vendrell, M., de Leau, E., Paterlini, A., Knox, K., Ross-Elliot, T., Reinders, A., Brockman, S.A., Ward, J., Oparka, K., 2015. Multispectral phloem-mobile probes: properties and applications. Plant Physiol. 167, 1211-1220.
2. Melnyk, C.W., Schuster, C., Leyser, O., Meyerowitz, E.M., 2015. A developmental framework for graft formation and vascular reconnection in *Arabidopsis thaliana*. Current Biol. 25, 1-13.
3. Yin, H., Yan, B., Sun, J., Jia, P.F., Zhang, Z.J., Yan, X.S., Chai, J., Ren, Z.Z., Zheng, G.C., Liu, H., 2012. Graft-union development: a delicate process that involves cell-cell communication between scion and stock for local auxin accumulation. J Exp Bot. 63, 4219-4232.

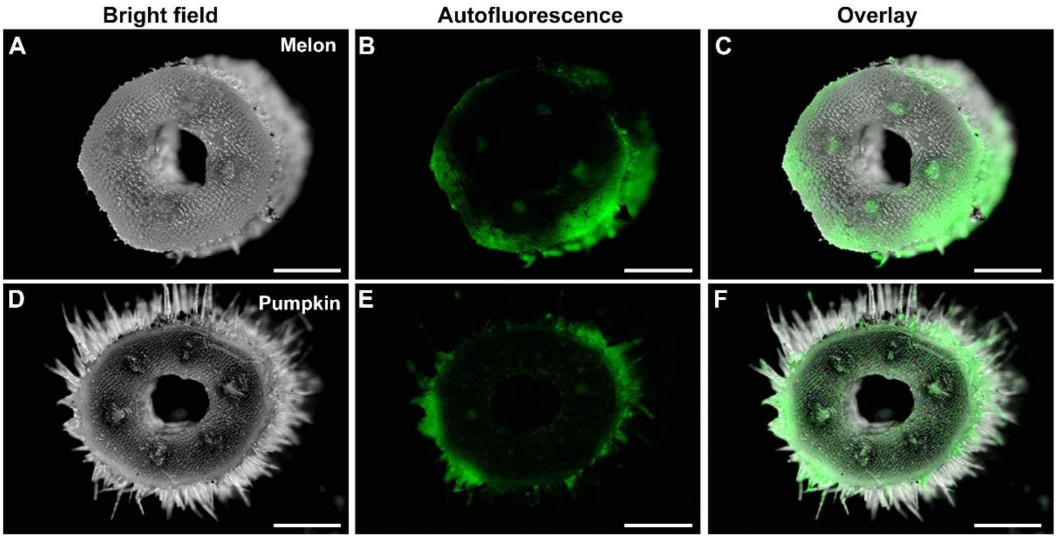


**Supplemental Figure S1. Melon and pumpkin stems showed strong auto-fluorescence under blue-light excitation.** Stem cross sections of 10-day-old (**A-C**) melon and (**D-F**) pumpkin seedlings were observed using a stereomicroscope. (A, D) Bright field. (B, E) Autofluorescence under blue-light excitation. (C, F) Overlay images of bright filed and fluorescence images. Bar: 1 mm.

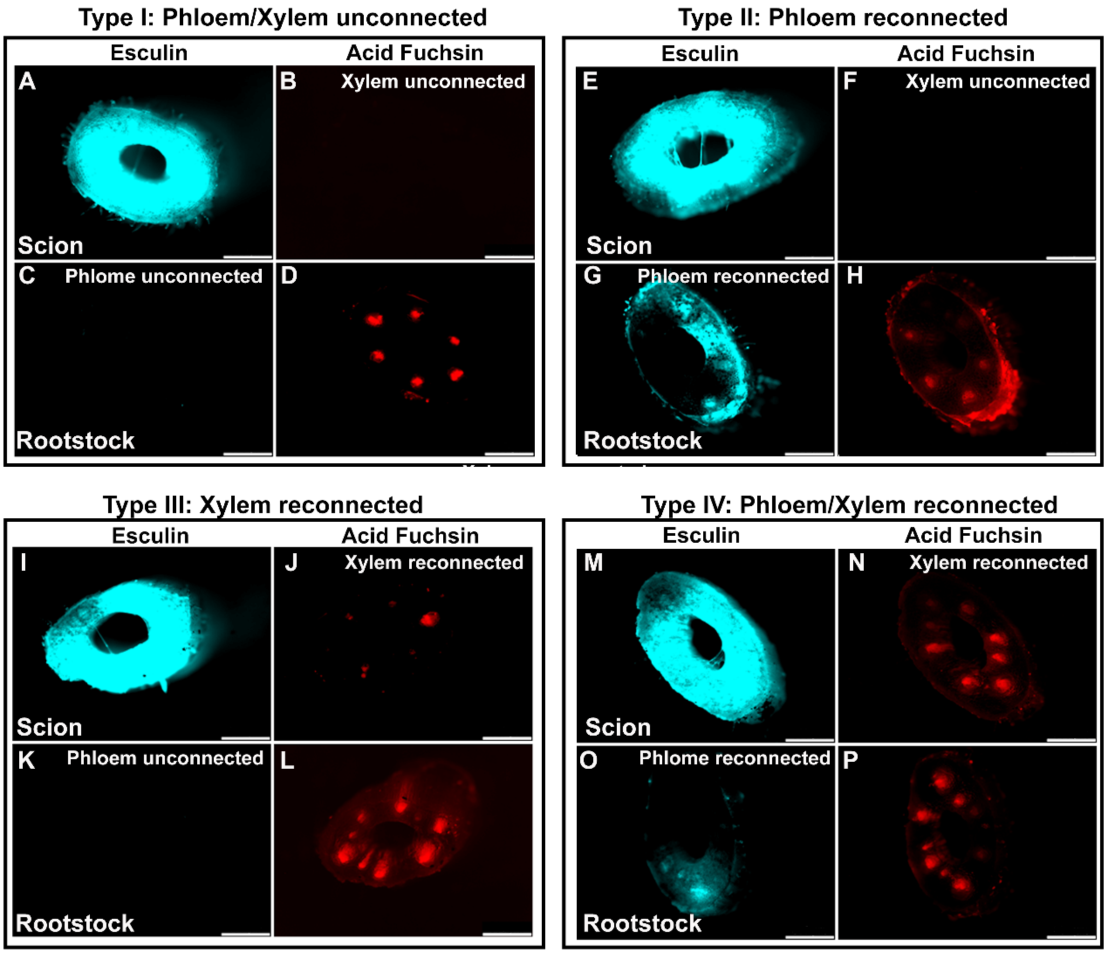


**Supplemental Figure S2. Cross sections from the scion and rootstock stems illustrating four different vascular connection types in grafted watermelon seedlings at 6 DAG.** To monitor phloem and xylem reconnections in grafted watermelon seedlings, fluorescent tracers including esculin and acid fuchsin were applied on the scion cotyledons and the rootstock roots, respectively. Stem sections from the scion and the rootstock of the same grafted seedlings were examined for the presences of esculin and acid fuchsin using fluorescent dissecting microscope. (**A-D**) Phloem and xylem were both unconnected. (**E-H**) Phloem was reconnected but xylem was unconnected. (**I-L**) Xylem was reconnected but phloem was unconnected. (**M-P**) Phloem and xylem were both reconnected. Bar: 1 mm.

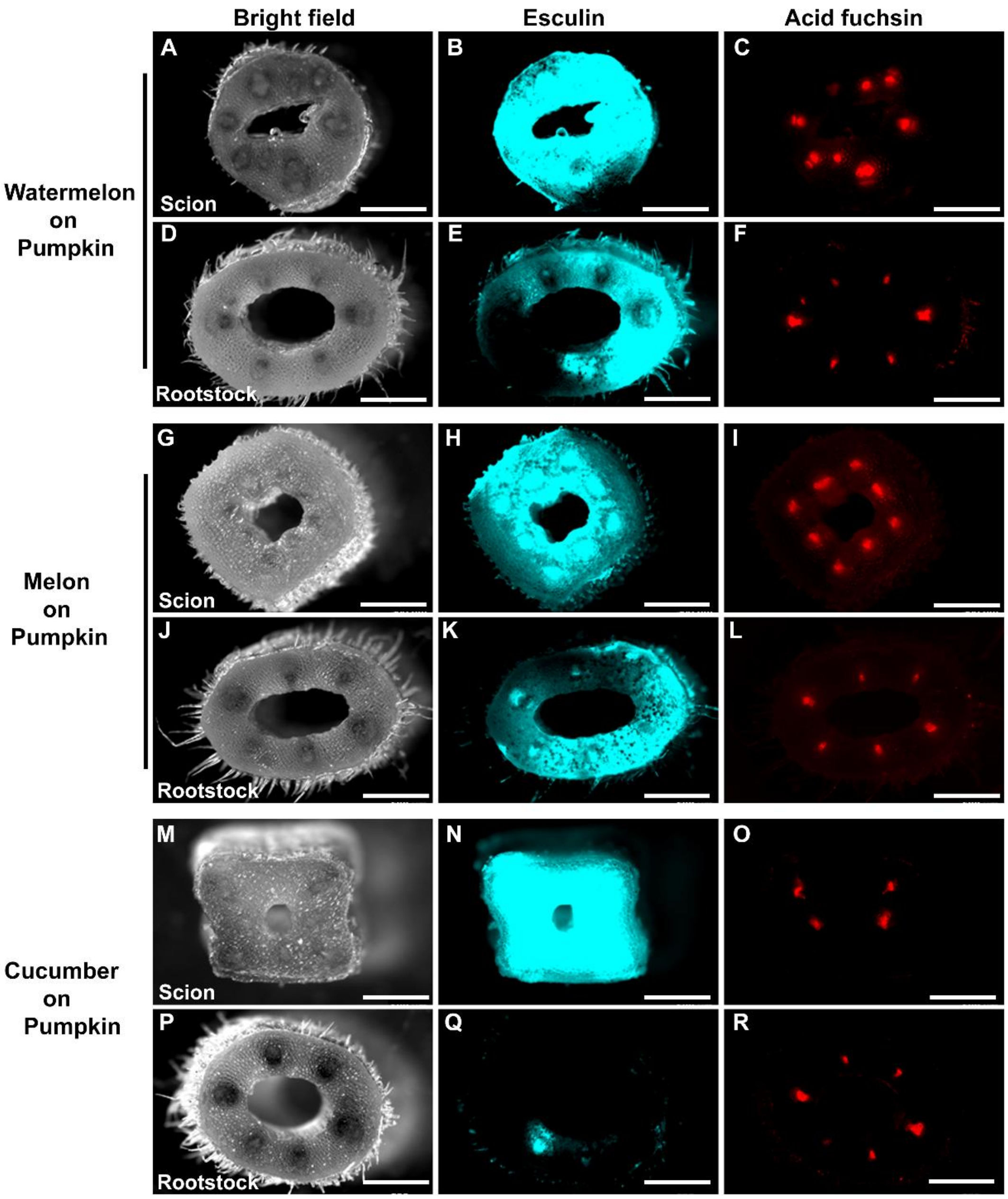


**Supplemental Figure S3. Application of esculin and acid fuchsin in heterograft combinations of watermelon, melon and cucumber onto pumpkin.** Esculin and acid fuchsin were applied on the scion cotyledons and the rootstock roots, respectively, to monitor reconnections of the phloem and the xylem tissues in the grafted seedlings of each heterograft combination. Scion stems from (**A-C**) Watermelon, (**G-I**) melon, and (**M-O**) cucumber those were grafted onto (**D-F**, **J-L**, **P-R**) the pumpkin rootstocks. Exogenously applied esculin could be transported from the scion cotyledon to the rootstock. Meanwhile, acid fuchsin fed to the root also could be loaded into the scion tissues via their reconnected xylem tissues. Bar: 1 mm.

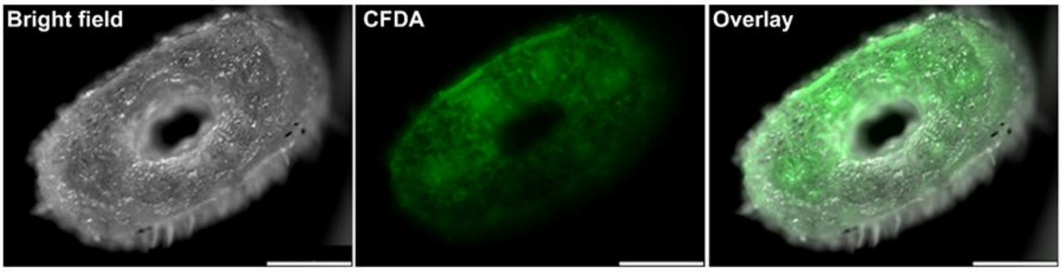


**Supplemental Figure S4. CFDA fluorescence observed in watermelon stem.** The presence of CFDA in the stem of 10-day-old watermelon seedling was detected under blue-light excitation using a stereomicroscope. Bar: 1 mm.
